# Phototrophic co-cultures from extreme environments: community structure and potential value for fundamental and applied research

**DOI:** 10.1101/427211

**Authors:** Claire Shaw, Charles Brooke, Erik Hawley, Morgan P. Connolly, Javier A. Garcia, Miranda Harmon-Smith, Nicole Shapiro, Michael Barton, Susannah G. Tringe, Tijana Glavina del Rio, David E. Culley, Richard Castenholz, Matthias Hess

**Affiliations:** Systems Microbiology & Natural Products Laboratory, University of California, Davis, CA; Bayer, Pittsburg, PA; Microbiology Graduate Group, University of California, Davis, CA; Biochemistry, Molecular, Cellular, and Developmental Biology Graduate Group, University of California, Davis, CA; Department of Energy, Joint Genome Institute, Berkeley, CA; Greenlight Biosciences, Medford, MA; University of Oregon, Eugene, OR

## Abstract

Cyanobacteria are found in most illuminated environments and are key players in global carbon and nitrogen cycling. Although significant efforts have been made to advance our understanding of this important phylum, still little is known about how members of the cyanobacteria affect and respond to changes in complex biological systems. This lack of knowledge is in part due to our dependence on pure cultures when determining the metabolism and function of a microorganism. In the work presented here we took advantage of the Culture Collection of Microorganisms from Extreme Environments (CCMEE), a collection of more than 1,000 publicly available photosynthetic co-cultures now maintained at the Pacific Northwest National Laboratory. To highlight some of their scientific potential, we selected 26 of these photosynthetic co-cultures from the CCMEE for 16S rRNA gene sequencing. We assessed if samples readily available from the CCMEE could be used to generate new insights into the role of microbial communities in global and local carbon and nitrogen cycling. Results from this work support the existing notion that culture depositories in general hold the potential to advance fundamental and applied research. If collections of co-cultures can be used to infer roles of the individual organisms remains to be seen and requires further investigation.

## INTRODUCTION

Cyanobacteria are photosynthetic prokaryotes that are found in the majority of illuminated habitats and are known to be some of the most morphologically diverse prokaryotes on our planet (Whitton and Potts, 2000). The global cyanobacterial biomass is estimated to total ~3×10^14^ g of carbon (Garcia-Pichel et al., 2003) and cyanobacteria may account for 20–30% of Earth’s primary photosynthetic productivity (Pisciotta et al., 2010). The efficient photosynthetic machinery of cyanobacteria has inspired growing interest in the utilization of cyanobacteria and cyanobacteria containing co-cultures in microbial fuel cells (Zhao et al., 2012; Gajda et al., 2015). In addition to having a global effect on the carbon cycle, cyanobacteria-mediated nitrogen fixation has been estimated to supply 20–50% of the nitrogen input in some marine environments (Karl et al., 1997). A detailed comprehension of cyanobacteria and their contribution to global carbon and nitrogen cycling is therefore indispensable for a multi-scalar and holistic understanding of these globally important nutrient cycles.

Besides their ecological relevance, cyanobacteria have potential applications in biotechnology: cyanobacteria facilitate the assimilation of carbon dioxide, a cheap and abundant substrate, to synthesize a variety of value-added compounds with industrial relevance (Al-Haj et al., 2016). Although monocultures have dominated in microbial biomanufacturing, controlled co-cultures have been recognized as valuable alternatives, due to their potential of reducing the risk of costly contaminations and in some cases enabling increasing product yield (Wang et al., 2015; Yen et al., 2015; Padmaperuma et al., 2018). Numerous cyanobacterial strains have been investigated for their potential to produce bioactive compounds, biofertilizer, biofuels, and bioplastics (Abed et al., 2009; Woo and Lee, 2017; Miao et al., 2018); and co-expression of non-cyanobacterial genes as well as co-cultivation of cyanobacteria with non-photosynthetic bacteria has resulted in self-sustained systems and improved desirable cyanobacterial phenotypes (de-Bashan et al., 2002; Subashchandrabose et al., 2011; Formighieri and Melis, 2016). Genes coding for enzymes capable of catalyzing reactions that result in unique products, such as modified trichamide, a cyclic peptide suggested to protect the bloom-forming *Trichodesmium erythraeum* against predation (Sudek et al., 2006), and prochlorosins, a family of lanthipeptides with diverse functions that are synthesized by various strains of *Prochlorococcus* and *Synechococcus* (Li et al., 2010; Cubillos-Ruiz et al., 2017), have been identified from cyanobacterial genomes (Zarzycki et al., 2013; Kleigrewe et al., 2016). It is very likely that *de novo* genome assembly from metagenomic data will facilitate the discovery of novel enzymes from cyanobacteria that are recalcitrant to current isolation and cultivation techniques. Although metagenome-derived genomes hold great potential to enhance our knowledge about genomic dark matter, improved techniques to isolate and enable axenic culturing of microorganisms that are currently considered as “unculturable”, as well as new genetic tools to study non-axenic cultures will be necessary in order to fully access the biotechnological potential of cyanobacteria.

Culture collections provide the possibility of preserving microbial isolates over extended periods of time without introducing significant genetic changes (McCluskey, 2017) and they facilitate open access to these isolates and their associated metadata (Boundy-Mills et al., 2015). Although culture collections hold enormous potential for capturing and preserving microbial biodiversity, there are numerous challenges in maintaining these biological depositories and the individual samples they contain. With recent advances in DNA sequencing technologies and the accessibility of 16S rRNA gene-based microbial community profiling, we are now well positioned to re-inventory, and standardize existing culture collections, which will be essential for preserving and cataloguing the planet’s microbial biodiversity.

To explore the potential of culture collections, specifically those that maintain samples of microbial co-cultures, we reexamined the biodiversity of 26 historical phototrophic samples from the Culture Collection of Microorganisms from Extreme Environments (CCMEE). While some of the samples, and their dominant phototrophs were studied previously using 16S rRNA profiling and morphological characterization (Camacho et al., 1996; Miller and Castenholz, 2000; Nadeau and Castenholz, 2000; Nadeau et al., 2001; Dillon et al., 2002; Dillon and Castenholz, 2003; Norris and Castenholz, 2006; Toplin et al., 2008) the diversity of the photosynthetic and non-photosynthetic organisms and the overall community assemblage of these co-cultures have not yet been characterized. To add further value to this study, we selected samples that originated from diverse extreme environments with distinct physical properties from across the globe, suggesting each co-culture would yield a unique microbial consortium. An enhanced understanding of the microbial diversity, even if affected by culturing conditions, of environmental co-cultures available through public culture collections, will contribute to a better understanding of global microbial biodiversity and demonstrate the utility of such culture collections.

## MATERIALS AND METHODS

### Sample collection & sample description

Co-cultures selected for this study are part of a larger culture collection and were collected from different locations (Table 1) between 1988 and 2002. Isolates were collected using sterile techniques, kept in the dark and stored on ice as soon as possible. Samples were transported to the laboratory where aliquots were prepared for cultivation and preservation at −80°C. For this study, co-cultures were selected from the CCMEE to cover a variety of geographical locations (Supplemental Figure S1) as well as a range of different ecosystems (Table 1). Due to the lack of a consistent terminology to describe the sampling sites, we categorized co-cultures according to the geographical location (e.g. Antarctica, Bermuda, Denmark, Mexico and Spain) and on a general description of ecosystem (i.e. creek, crust, freshwater, hot spring, marine, saline pond, terrestrial, travertine, and tree bark) from where the co-cultures were collected. In addition, we used the growth medium and temperature (i.e. 12°C, 23°C, 40°C, 45°C, 55°C) at which available co-cultures have been maintained historically in the CCMEE to categorize the co-cultures used in this study.

**Table 1:** Summary of photosynthetic co-cultures for which 16S rRNA gene profiles were generated.

| FECB ID | Growth Temperature [°C] | Sample Location (country) | Sample Location (region) | Habitat |
| --- | --- | --- | --- | --- |
| FECB1 | 12 | Antarctica | McMurdo Ice Shelf; Bratina Island | Pond (saline) |
| FECB2 | 12 | Antarctica | McMurdo Ice Shelf; Bratina Island | Pond (freshwater) |
| FECB3 | 12 | Antarctica | McMurdo Ice Shelf; Bratina Island | Pond (brackish) |
| FECB4 | 23 | Spain | Lake Arcas | Lake (freshwater) |
| FECB5 | 23 | Spain | Lake Arcas | Lake (freshwater) |
| FECB6 | 23 | United States | Yellowstone National Park; Pott's Basin | Hot Spring |
| FECB10 | 23 | Mexico | Viscaino Desert | Terrestrial (epilithic) |
| FECB12 | 23 | United States | University of Oregon; Eugene, Oregon | Terrestrial (concrete) |
| FECB14 | 23 | United States | Yellowstone National Park; Pott's Geyser | Hot Spring |
| FECB15 | 23 | United States | Yellowstone National Park; Rabbit Creek | Warm Creek |
| FECB17 | 23 | United States | Yellowstone National Park; Rabbit Creek | Warm Creek |
| FECB19 | 23 | United States | Yellowstone National Park; Shoshone Geyser Basin | Hot Spring |
| FECB21 | 23 | United States | Yellowstone National Park; Pott's Geyser | Hot Spring |
| FECB22 | 23 | United States | Hawaii | Terrestrial (tree trunk) |
| FECB24 | 23 | Antarctica | McMurdo Ice Shelf; Bratina Island | Pond (freshwater) |
| FECB26 | 23 | Bermuda | Sommerset | Terrestrial (wooden fence) |
| FECB28 | 23 | Antarctica | McMurdo Ice Shelf; Bratina Island | Pond (saline) |
| FECB30 | 23 | Denmark | Limfjord Shallows, Limfjord | Marine |
| FECB32 | 23 | United States | Yellowstone National Park; Mammoth; Narrow Gauge | Travertine (endolithic) |
| FECB34 | 23 | United States | Yellowstone National Park; Lower Geyser Basin; Great Fountain | Crust |
| FECB36 | 23 | United States | Yellow Stone National Park; Lower Geyser Basin; Sentinel Spring | Hot Spring (silica crust) |
| FECB38 | 23 | United States | Yellowstone National Park; Lower Geyser Basin; Mushroom Spring | Crust |
| FECB52 | 40 | United States | Yellowstone National Park; Norris Geyser Basin | Terrestrial (acid crust) |
| FECB53 | 40 | United States | Yellowstone National Park; Sylvan Springs | Hot Spring (acid pool) |
| FECB58 | 45 | United States | Hunter's Hot Spring, Oregon | Hot Spring |
| FECB68 | 55 | United States | Hunter's Hot Spring, Oregon | Hot Spring |

FECB1 (CCMEE ID 5011) and FECB3 (CCMEE ID 5034) were collected from saline and brackish melt ponds and were dominated by phototrophic cyanobacteria classified as *Oscillatoria* sp. (Nadeau et al., 2001). FECB2 (CCMEE ID 5019) was collected from a freshwater pond and was phylogenetically uncharacterized prior to our efforts. FECB4 (CCMEE ID 5047; AP1) and FECB5 (CCMEE ID 5049; AO21) were also isolated from freshwater and the dominant photosynthetic organisms within these samples were classified by 16S rRNA sequence analysis as being related to *Pseudanabaena limnetica* and *Oscillatoria* cf. *tenuis*, respectively (Camacho et al., 1996). FECB6 (CCMEE ID 5051), FECB14 (CCMEE ID 5093; WT-97 Cal), FECB15 (CCMEE ID 5083), and FECB19 (CCMEE ID 5091; Y-97) were collected from diverse hot springs within Yellowstone National Park (YNP) (Table 1). FECB10 (CCMEE ID 5056; M88-VD (1)) was collected as epiliths (Dillon et al., 2002). FECB17 (CCMEE ID 5085; RC-97 Cal) and FECB36 (CCMEE ID 6076) were isolated from Rabbit Creek and a crust in the Sentinel Spring Meadows in YNP respectively and dominant phototrophs of these co-cultures were characterized previously as *Calothrix* spp. (Dillon and Castenholz, 2003). FECB22 (CCMEE ID 5097; HW-91) and FECB26 (CCMEE ID 5099; B77-scy,j,) were collected from a tree trunk and a wooded fence respectively. FECB24 (CCMEE ID 5098; AN-90) was collected from a shallow melt pond (~10 m^2^) in the Victoria Valley, Antarctica, whereas FECB28 (CCMEE ID 5102) was collected from a saline melt pond on Bratina Island, Antarctica (Nadeau and Castenholz, 2000). FECB32 (CCMEE ID 6031), FECB34 (CCMEE ID 6069) and FECB38 (CCMEE ID 6083) were endoliths collected from subsurface (1-5 mm depths) travertine deposits in YNP (Norris and Castenholz, 2006). FECB53 (CCMEE ID 5610) was collected from Sylvan Springs in YNP. Temperature and pH at FECB53’s sampling site were determined to be 40 °C and pH4, conditions which are considered to be too harsh to actively support growth of cyanobacteria, and Toplin et al. reported the thermo-acidophilic red algae *Cyanidioschyzon* as a highly abundant phototropic strain in this sample (Toplin et al., 2008). FECB58 (CCMEE ID 5216; OH-9-45C) and FECB68 (CCMEE ID 5240; OH-2-55C) were collected from Hunter’s Hot Spring in Oregon and in 2000 Miller and Castenholz reported the isolation of several thermophilic clones belonging to the genus *Synechococcus* from these samples (Miller and Castenholz, 2000).

### Growth of co-cultures

To obtain sufficient biomass for subsequent DNA analysis, 100 μL of each co-culture were transferred to 25 mL of sterile BG11 media (Allen and Stanier, 1968). For FECB52 and FECB53 BG11 was substituted by Cyanidium medium (Castenholz, 1981). Co-cultures were subjected to a 12 hr diurnal light/dark cycle while grown over 28 days at a temperature somewhat similar to the temperature that was measured at the location where to sample was located. Growth temperature for each sample is indicated in Table 1.

### DNA extraction and 16S rRNA gene amplification

Total microbial DNA was extracted from 500 μL of each photosynthetic co-culture using the FastDNA SPIN Kit for Soil (MP Biomedical, Solon, OH) according to the manufacturer’s instructions. Extracted DNA was quantified via fluorescence (Qubit; Thermo Scientific, USA) and the hypervariable V4 region of the 16S rRNA gene was amplified from extracted DNA using the primer set 515F/805R (515F: 5’-GTGCCAGCMGCCGCGGTAA-3’ and 805R: 5’-GGACTACHVGGGTWTCTAAT-3’). The forward primer included an 11 bp barcode to allow multiplexing of samples during sequencing. The barcode sequence for each sample is listed in Supplemental Table S1. Subsequent PCR reactions were performed using the 5PRIME HotMasterMix amplification mix (QIAGEN, Beverly, MA) with the following PCR conditions: initial denaturation for 90 sec at 94°C, followed by 30 amplification cycles (45 sec at 94°C, 60 sec at 60°C, and 90 sec at 72°C) followed by a final extension step of 72°C for 10 min. Amplification products were cooled to 4°C. Samples were sequenced at the Department of Energy’s Joint Genome Institute (JGI; http://www.jgi.doe.gov) according to JGI’s standard operating procedure using Illumina’s MiSeq platform and v3 chemistry.

### Sequence data analysis

Raw sequencing data were downloaded from the JGI’s Genome Portal (http://genome.jgi.doe.gov/) under the project ID 1032475. Data were decompressed and de-interleaved using the 7-zip software (www.7-zip.org) and an in-house script, respectively. De-interleaved files were subsequently processed using MOTHUR version 1.38.1 (Schloss et al., 2009; Kozich et al., 2013). Paired-end reads were combined using the *make.contigs* command. Sequences with ambiguous base calls and sequences longer than 325 bp were removed using *screen.seqs*. Duplicate sequences were merged using *unique.seqs*, and the resulting unique sequences were aligned to the V4 region of the SILVA database (v123) (Quast et al., 2013). Chimeras were removed using UCHIME (Edgar et al., 2011) and quality filtered sequences were taxonomically classified at 80% confidence to the GreenGenes reference taxonomy (release gg_13_5_99) (McDonald et al., 2012). Non-prokaryotic sequences were removed and the *dist.seqs* command was used to calculate pairwise distances between the aligned sequences. The resulting pairwise distance matrix was used to cluster sequences into operational taxonomic units (OTUs) with a 97% sequence identity cut-off using UCLUST (Edgar, 2010). The most abundant sequence of each OTU was picked as the representative sequence. OTUs were taxonomically classified using the *classify.otu* command using the GreenGenes reference taxonomy (release gg_13_5_99). Shannon and Simpson estimators were calculated in MOTHUR (Schloss et al., 2009).

In order to visualize the overall compositional differences between the co-cultures, an uncorrected pairwise distance matrix was generated using the *dist.seqs* command in MOTHUR and a tree was generated using *Clearcut* (version 1.0.9) (Evans et al., 2006). A cladogram from the resulting tree file was constructed and visualized using iTOL (Letunic and Bork, 2016). Cluster designations were assigned at a branch length of 0.05, with branch length indicating the (number of differences / overall length of branches) between two samples. Samples whose branches split at a distance >0.05 were considered as part of the same cluster (Figure 1).

**Figure 1:**
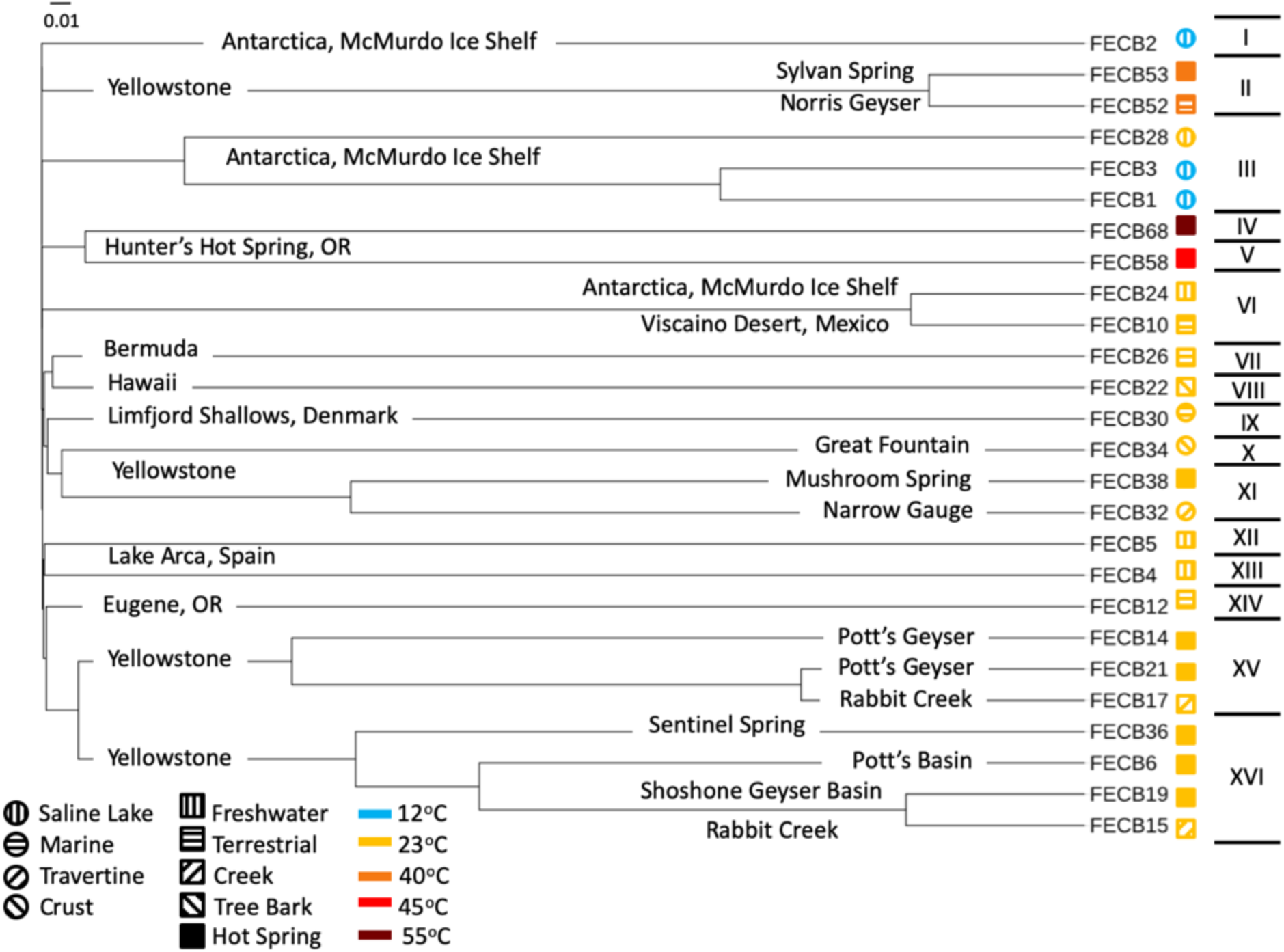
Cladogram of 16S rRNA based community composition of co-cultures under investigation. FECB identifier (sample ID) is provided for each co-culture. Sample location is indicated on the corresponding branch. Roman numerals on the right indicate the clusters identified at a branch cutoff of 0.05. Symbols (i.e. circles and squares) next to sample ID indicate habitat type and color indicates the temperatures at which sample was historically maintained in the CCMEE. Branch length indicates (number of differences / overall length of branches) between two samples.

### Availability of data and material

Co-cultures subject to this study are publicly available through the CCMEE and the UTEX Culture Collection of Algae at the University of Texas at Austin upon request using the corresponding FECB ID (Table 1). Co-cultures can also be obtained from the Hess Lab at UC Davis. Sequences generated during this project have been deposited and are publicly available at NCBI’s SRA under the BioProject ID PRJNA401502. All other data is included in this published article and its supplementary information files.

## RESULTS & DISCUSSION

A total of 3,357,905 raw reads (mean (SD) = 129,150 (±15,845) reads per sample) were generated from the V4 region of the 16S rRNA gene (Table 2). Quality filtering removed ~3.8% (±0.57%) of the raw reads from each sample due to insufficient quality. The remaining reads were assigned to a total of 5,785 distinct Operational Taxonomic Units (OTUs) based on 97% sequence identity (Table S2).

**Table 2:** Read statistics and Diversity Index for co-cultures investigated in this study.

| Sample ID | Total Raw Reads | Total Quality Filtered Reads | Total OTUs Observed | OTUs Recruiting >0.1% of Reads | % Reads Represented by OTUs Recruiting >0.1% of Reads | % Reads Recruited by Dominant OTU | Inverse Simpson Index | Shannon Index |
| --- | --- | --- | --- | --- | --- | --- | --- | --- |
| FECB1 | 154,251 | 139,666 | 349 | 13 | 99.6% | 47% | 3.23 | 1.49 |
| FECB2 | 128,172 | 116,166 | 197 | 3 | 99.8% | 51% | 2.04 | 0.77 |
| FECB3 | 139,819 | 128,879 | 274 | 8 | 99.6% | 80% | 1.52 | 0.76 |
| FECB4 | 94,960 | 84,983 | 288 | 17 | 99.4% | 44% | 3.92 | 1.77 |
| FECB5 | 122,414 | 108,882 | 401 | 18 | 99.4% | 40% | 4.63 | 2.00 |
| FECB6 | 153,355 | 136,199 | 549 | 28 | 99.4% | 29% | 6.5 | 2.39 |
| FECB10 | 114,324 | 102,534 | 299 | 10 | 99.7% | 72% | 1.78 | 0.96 |
| FECB12 | 151,629 | 141,953 | 335 | 19 | 99.5% | 72% | 1.87 | 1.23 |
| FECB14 | 145,839 | 98,588 | 206 | 4 | 99.6% | 67% | 1.83 | 0.75 |
| FECB15 | 105,256 | 96,030 | 278 | 9 | 99.6% | 63% | 2.18 | 1.14 |
| FECB17 | 109,474 | 97,391 | 317 | 7 | 99.5% | 59% | 2.47 | 1.33 |
| FECB19 | 118,003 | 103,477 | 300 | 10 | 99.5% | 47% | 3.27 | 1.50 |
| FECB21 | 127,371 | 113,281 | 401 | 15 | 99.2% | 58% | 2.38 | 1.27 |
| FECB22 | 129,934 | 117,651 | 399 | 23 | 99.4% | 40% | 4.46 | 2.08 |
| FECB24 | 115,275 | 93,362 | 265 | 12 | 99.9% | 65% | 2.12 | 1.17 |
| FECB26 | 121,303 | 103,597 | 265 | 16 | 99.5% | 63% | 2.29 | 1.32 |
| FECB28 | 152,974 | 134,318 | 332 | 10 | 99.4% | 37% | 3.22 | 1.36 |
| FECB30 | 140,681 | 118,256 | 546 | 22 | 98.7% | 33% | 4.61 | 2.00 |
| FECB32 | 116,889 | 72,130 | 336 | 29 | 98.3% | 19% | 9.24 | 2.68 |
| FECB34 | 138,723 | 125,710 | 288 | 14 | 99.9% | 76% | 1.69 | 1.11 |
| FECB36 | 137,395 | 124,885 | 360 | 12 | 99.6% | 52% | 2.38 | 1.16 |
| FECB38 | 139,001 | 124,379 | 479 | 17 | 99.1% | 32% | 5.22 | 2.01 |
| FECB52 | 117,261 | 75,433 | 367 | 7 | 99.9% | 71% | 1.85 | 1.02 |
| FECB53 | 119,380 | 86,565 | 303 | 5 | 99.1% | 91% | 1.21 | 0.44 |
| FECB58 | 135,683 | 124,119 | 351 | 13 | 99.3% | 38% | 3.65 | 1.61 |
| FECB68 | 128,539 | 114,032 | 402 | 11 | 99.7% | 26% | 5.27 | 1.90 |
| **Total** | 3,357,905 | 2,882,466 | N/A | N/A | N/A | N/A | N/A | N/A |
| **Average** | 129,150 | 110,864 | 342 | 14 | 99% | 51% | N/A | N/A |
| **Stdev** | 15,845 | 19,275 | 87.2 | 6 | 0% | 17% | N/A | N/A |

To estimate the microbial diversity within each sample, rarefaction analyses were performed (Supplemental Figure S2) and diversity indices were calculated (Table 2). The Inverse Simpson index (Simpson, 1949; Morris et al., 2014) of the samples ranged between 1.21 and 9.24 with the lowest and highest indices calculated for FECB53 and FECB32 respectively (Table 2). Not surprisingly, the diversity in the co-cultures under investigation appeared to be negatively correlated with the proportion of reads recruited by the dominant OTU of each sample (Pearson *r* =-0.8806; *p* <0.01). Although samples ranked slightly differently based on their diversity, when diversity was calculated using the Shannon index (Kim et al., 2017), the overall trend remained the same (Table 2).

### The McMurdo Dry Valley Lake System, a physically highly stable lacustrine system

The McMurdo Dry Valley (MDV) is one of the most extreme deserts on Earth, and although the importance of microbial communities for biogeochemical cycles of this region is widely accepted, the microbial ecology of the MDV remains poorly understood (Chan et al., 2013). FECB3, originating from a brackish pond on Bratina Island, was dominated by OTU000003, which recruited 80.3% of all reads (Supplemental Table S2). OTU000003 was classified as the cyanobacterium *Phormidium pseudopriestleyi*, previously reported to dominate microbial mats of the anoxic zone of Lake Fryxell, Antarctica (Jungblut et al., 2015). The second and third most abundant OTUs in FECB3 were OTU000015 and OTU000061, respectively (Supplemental Table S2). Both OTU000015 and OTU000061 were classified as Rhodobacteriaceae and recruited 9.2% and 8.2% of the reads generated for FECB3. Whereas a taxonomic classification of OTU000015 was not possible beyond the family level, OTU000061 was classified as *Paracoccus marcusii*, a Gram-negative organism that displays a bright orange color due to the synthesis of carotenoids such as astaxanthin (Harker et al., 1998).

While the microbial ecology of melt ponds and lakes in the MDV, habitats covered year-round with an ice sheet, have been studied in great detail, most of the insights regarding the microbial community assemblage in these waters are based primarily on visual observations via microscopy (Jungblut et al., 2015). Molecular data, like those presented here, will be of great value to extend our knowledge framework of the microbial ecology of this unique ecosystem.

### Ubiquity of Cyanobacteria and Proteobacteria within photosynthetic co-cultures

While the microbial communities of the co-cultures under investigation varied greatly, cyanobacteria and proteobacteria co-occurred in all 26 of the community assemblages. Community composition analysis revealed that each of the co-cultures contained at least one OTU (mean (SD) = 2 (±1.23)) that recruited >0.1% of the co-culture specific reads and that was classified as *Cyanobacteria* (Table 3). The only other phylum present in each of the individual 26 co-cultures and represented by at least one OTU recruiting >0.1% of the reads was the *Proteobacteria* phylum (Table 3). In contrast, only three samples, namely FECB5, FECB30 and FECB68, contained OTUs that recruited >0.1% of the sample specific reads and that could not be classified at the phylum level or at a higher taxonomic resolution (Table 3). It is possible that the relatively high abundance of non-classified phyla might contribute to the separation of these samples into distinct clusters (i.e. cluster XII, IX, and IV; Figure 1). In addition to their ubiquity, *Cyanobacteria* and *Proteobacteria* also recruited the majority of the reads in all but four (i.e. FECB2, FECB12, FECB58, and FECB68) of the samples under investigation (Figure 2 and Supplemental Table S3). In FECB2 and FECB12 the majority of the reads were recruited by OTUs classified as members of the phylum *Bacteroidetes* (recruiting 50.6% and 72% of the reads respectively), whereas within FECB58 and FECB68, *Armatimonadetes* (38.3%) and *Chloroflexi* (25.9%) were identified as the most abundant phyla (Figure 2 and Supplemental Table S3). The fact that these samples were dominated by phyla other than the *Cyanobacteria* or *Proteobacteria* may also help to explain why these samples form distinct clusters (cluster I, XIV and V, IV respectively; Figure 1).

**Table 3:** Count and phylogenetic classification of identified OTUs at the phylum level. Only OTUs recruiting >0.1% of the co-culture specific reads are shown.

| Phylum | FECB 1 | FECB 2 | FECB 3 | FECB 4 | FECB 5 | FECB 6 | FECB 10 | FECB 12 | FECB 14 | FECB 15 | FECB 17 | FECB 19 | FECB 21 | FECB 22 | FECB 24 | FECB 26 | FECB 28 | FECB 30 | FECB 32 | FECB 34 | FECB 36 | FECB 38 | FECB 52 | FECB 53 | FECB 58 | FECB 68 |
| --- | --- | --- | --- | --- | --- | --- | --- | --- | --- | --- | --- | --- | --- | --- | --- | --- | --- | --- | --- | --- | --- | --- | --- | --- | --- | --- |
| Cyanobacteria | 2 | 1 | 2 | 1 | 1 | 3 | 1 | 2 | 2 | 2 | 1 | 2 | 6 | 5 | 1 | 1 | 1 | 2 | 3 | 1 | 2 | 2 | 3 | 1 | 1 | 2 |
| Proteobacteria | 10 | 1 | 5 | 13 | 11 | 18 | 7 | 12 | 2 | 7 | 5 | 5 | 6 | 16 | 10 | 14 | 8 | 16 | 20 | 11 | 9 | 14 | 3 | 2 | 6 | 2 |
| Actinobacteria | 1 | 0 | 1 | 3 | 1 | 1 | 2 | 3 | 0 | 0 | 1 | 2 | 1 | 2 | 0 | 0 | 1 | 0 | 3 | 0 | 1 | 0 | 0 | 2 | 0 | 0 |
| Bacteroidetes | 0 | 1 | 0 | 0 | 1 | 6 | 0 | 1 | 0 | 0 | 0 | 0 | 2 | 0 | 1 | 1 | 0 | 1 | 1 | 1 | 0 | 1 | 0 | 0 | 1 | 0 |
| Armatimonadetes | 0 | 0 | 0 | 0 | 1 | 0 | 0 | 1 | 0 | 0 | 0 | 0 | 0 | 0 | 0 | 0 | 0 | 0 | 0 | 0 | 0 | 0 | 0 | 0 | 1 | 1 |
| Unclassified Bacteria | 0 | 0 | 0 | 0 | 1 | 0 | 0 | 0 | 0 | 0 | 0 | 0 | 0 | 0 | 0 | 0 | 0 | 1 | 0 | 0 | 0 | 0 | 0 | 0 | 0 | 2 |
| Firmicutes | 0 | 0 | 0 | 0 | 0 | 0 | 0 | 0 | 0 | 0 | 0 | 0 | 0 | 0 | 0 | 0 | 0 | 0 | 0 | 1 | 0 | 0 | 1 | 0 | 1 | 1 |
| Thermi | 0 | 0 | 0 | 0 | 1 | 0 | 0 | 0 | 0 | 0 | 0 | 0 | 0 | 0 | 0 | 0 | 0 | 0 | 0 | 0 | 0 | 0 | 0 | 0 | 0 | 1 |
| Chloroflexi | 0 | 0 | 0 | 0 | 1 | 0 | 0 | 0 | 0 | 0 | 0 | 0 | 0 | 0 | 0 | 0 | 0 | 0 | 0 | 0 | 0 | 0 | 0 | 0 | 2 | 2 |
| Acidobacteria | 0 | 0 | 0 | 0 | 0 | 0 | 0 | 0 | 0 | 0 | 0 | 0 | 0 | 0 | 0 | 0 | 0 | 0 | 1 | 0 | 0 | 0 | 0 | 0 | 1 | 0 |
| Planctomycetes | 0 | 0 | 0 | 0 | 0 | 0 | 0 | 0 | 0 | 0 | 0 | 1 | 0 | 0 | 0 | 0 | 0 | 2 | 1 | 0 | 0 | 0 | 0 | 0 | 0 | 0 |
| **Total OTU Count** | 13 | 3 | 8 | 17 | 18 | 28 | 10 | 19 | 4 | 9 | 7 | 10 | 15 | 23 | 12 | 16 | 10 | 22 | 29 | 14 | 12 | 17 | 7 | 5 | 13 | 11 |

**Figure 2:**
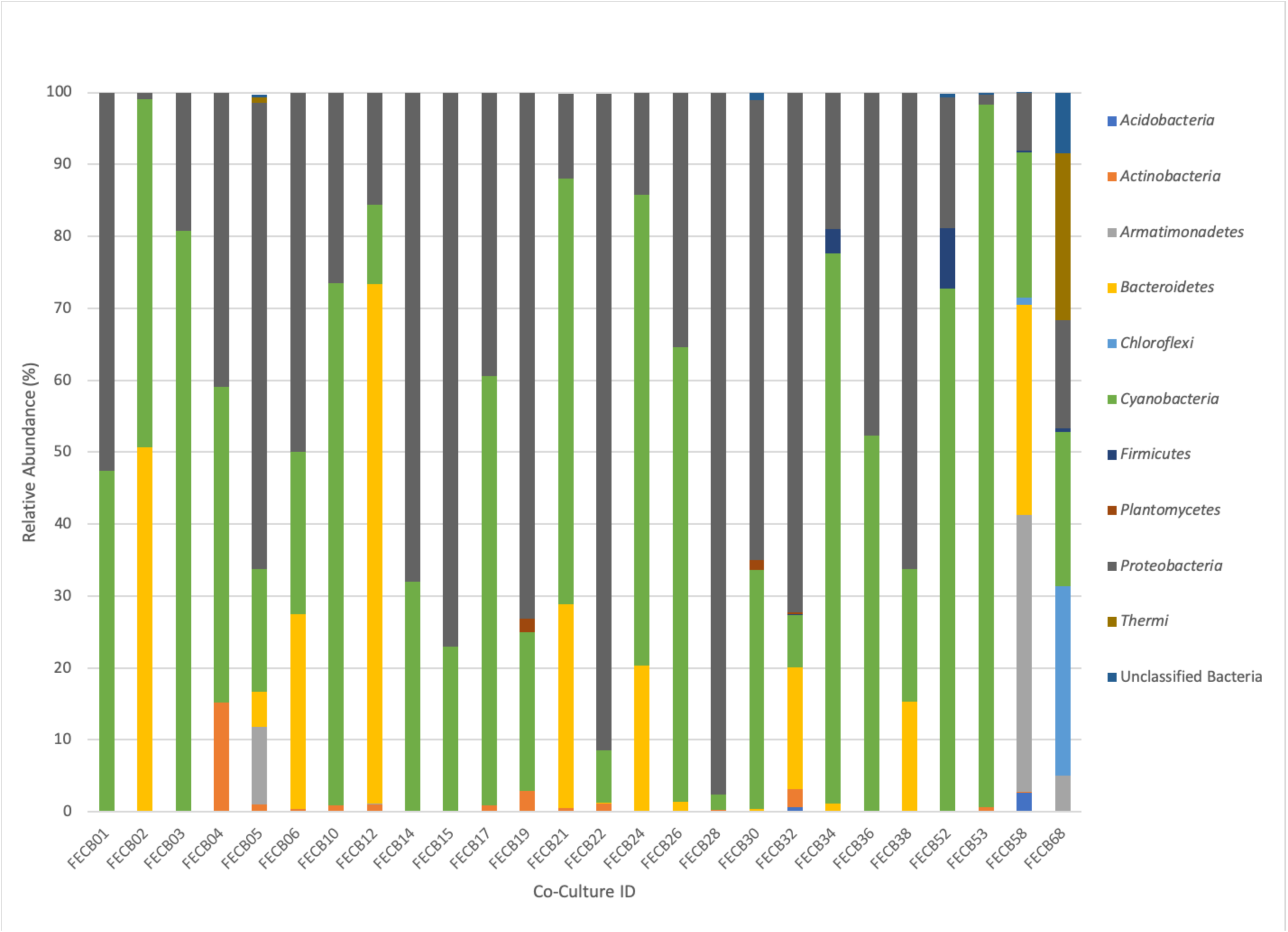
Relative abundance of phyla associated with phototrophic co-cultures. 16S rRNA based community profile. Only phyla recruiting >0.1% of the reads in at least one of the co-cultures are shown.

### Firmicutes dominate photosynthetic co-cultures from hot springs

*Firmicutes* abundances calculated for co-cultures from hot spring samples were higher compared to those calculated for co-cultures from other environments. OTUs assigned to the *Firmicutes* phylum were detected above the applied cut-off level of 0.1% in only five of the twenty-six co-cultures under investigation (Table 3). Interestingly, these samples (i.e. FECB32, FECB34, FECB52, FECB58 and FECB68) are co-cultures collected from hot springs or from deposits within hot springs, with FECB52, FECB58 and FECB68 being maintained in culture at temperatures >40°C. OTU000073 (classified as *Alicyclobacillus tolerans*), OTU00082 (classified as members of the genus *Paenibacillus*), OTU000154 (classified as *Geobacillus vulcani*), and OTU000158 (classified as a member of the *Bacillaceae* family) recruited 5.9%, 3.4%, 0.5% and 0.4% of the reads generated from FECB52, FECB34, FECB68 and FECB58 respectively (Supplemental Table S2). *Alicyclobacillus tolerans* and *Geobacillus vulcani* have been described previously as aerobic spore-forming thermophiles and have been isolated from lead–zinc ores (Karavaiko et al., 2005) and hot springs (Nazina et al., 2004) located in Russia, respectively. Members of the genus *Paenibacillus* have been isolated from a wide variety of environments and some *Paenibacillus* species have been found to promote crop growth directly via biological nitrogen fixation, phosphate solubilization, production of the phytohormone indole-3-acetic acid; and they have been identified as a potential source of novel antimicrobial agents (Grady et al., 2016). Although it is difficult to make a reliable prediction of the metabolic capacities of the organism associated with OTU000082 solely based on 16S rRNA data, it is certainly possible that this organism might possess the ability to promote or inhibit plant and microbial growth respectively.

### Photosynthetic co-cultures from Antarctica and YNP to study adaptation to increased radiation, low temperatures and oligotrophic growth conditions

Microbial adaptation to extreme environments and the molecular framework that enable microorganisms to survive and thrive in the presence of increased rates of radiation, low temperatures and in the absence of nutrients has fascinated the scientific community for decades and remains poorly understood. In an attempt to provide a better basis of the taxonomic make-up of co-cultures that were collected from ecosystems that are characterized by these extremes, we included co-cultures from Antarctica and YNP in this study (Table 1). OTU-based comparison of Antarctica and YNP co-cultures revealed between 197 (FECB2) and 549 (FECB6) distinct OTUs (mean (SD) = 342 (±87.2) OTUs) based on 97% sequence similarity (Table 2). The number of OTUs that recruited >0.1% of all reads ranged from 3 to 29 OTUs, with FECB2 and FECB32 having the lowest and highest OTU count respectively (Table 2). FECB2 was dominated by an OTU classified as *Hymenobacter*, which recruited all *Bacteroidetes-specific* reads generated from this sample (Tables 3 & 4). The genus *Hymenobacter* contains several pigmented bacteria that have been isolated from Antarctica and have been reported to possess increased resistance to radiation (Oh et al., 2016; Marizcurrena et al., 2017), which might explain their increased abundance in FECB2, a co-culture isolated from an environment known to possess increased levels of UV radiation. Taking this into consideration, FECB2 and its individual community members could be a potential target for future studies to enhance our understanding of processes that infer resistance to radiation and DNA damage. The second most abundant OTU in FECB2, recruiting 48% of the generated samples, was classified as *Phormidium* sp (Supplemental Table S2), a cyanobacterial genus that has been reported to dominate aquatic microbial mats from Antarctica (Jungblut et al., 2005; Strunecky et al., 2012). Representative isolates from this genus have been proposed previously as cost-effective options for industrial carotenoid production(Shukla and Kashyap, 2003), suggesting that FECB2 may hold the potential for this process.

**Table 4.**
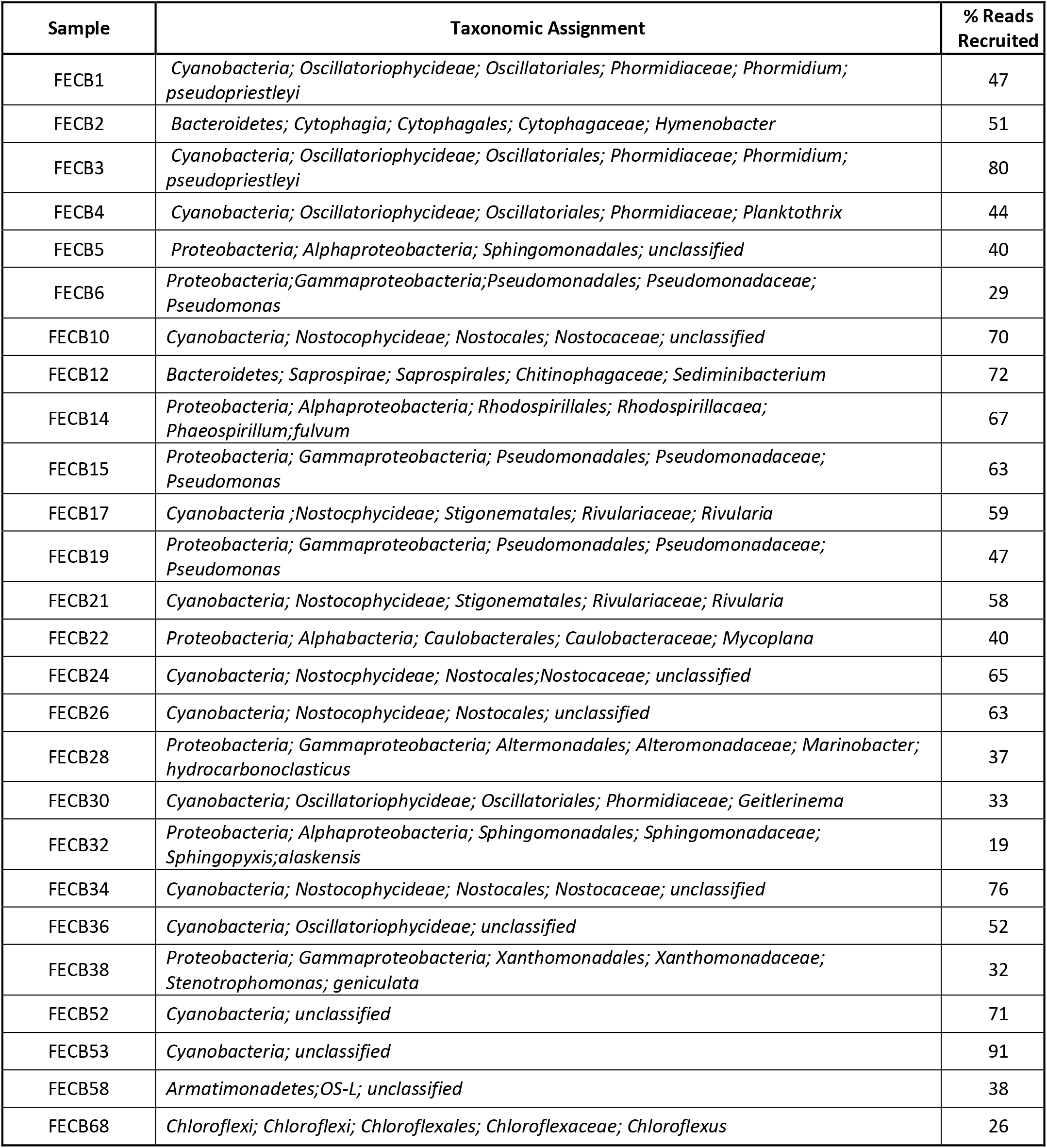
Taxonomy relative abundance of dominant OTU identified in each co-culture.

FECB32 is a mixed culture isolated from an ancient travertine at Mammoth in YNP. Our analysis indicated that FECB32 contained 29 OTUs that each accounted for >0.1% of the reads generated (Table 2). Fifteen of these OTUs recruited >1% of all reads and 4 OTUs collectively accounted for ~60% of the reads generated from this co-culture (Supplemental Table S4). These 4 OTUs were classified as *Sphingopyxis alaskensis, Chelativorans sp*. and as members of the *Chitinophagaceae* and *Comamonadaceae families*, recruiting ~19%, 13%, 17%, and 11% of the reads respectively (Supplemental Tables S2 & S4). *Sphingopyxis alaskensis* is a Gram-negative bacterium found in relatively high abundance in oligotrophic regions of the ocean (Vancanneyt et al., 2001; Cavicchioli et al., 2003) and it has been studied in great detail as a model system for marine bacteria, specifically to understand microbial adaptation to cold or oligotrophic environments (Lauro et al., 2009; Ting et al., 2010). The *Chitinophagaceae* family contains a wide phylogenetic diversity with many of its members being mesophilic. However, *Chitinophagaceae* have been reported to grow optimally at temperatures of 55°C and higher (Anders et al., 2014; Hanada et al., 2014).

### Photosynthetic co-cultures containing the deep-branching candidate phylum Melainabacteria

Extreme environments similar to those on early Earth are often proposed to hold critical information about the historical progression of life on our planet and a niche that encompasses those physical stresses is the endolithic environment of rocks (Norris and Castenholz, 2006). Phylogenetic analysis of the heterotrophic population associated with FECB32, which was isolated from travertine deposited by hot springs in YNP, found that sequences from MLE-12 (OTU000109) recruited ~2% of the sample specific sequences (Supplemental Table S2). This rendered MLE-12, previously assigned to the deep-branching candidate phylum *Melainabacteria* (Di Rienzi et al., 2013), as the eleventh most abundant organism in this photosynthetic co-culture. It has been proposed previously that *Melainabacteria*, which is commonly found in aquatic habitats, separated from the cyanobacteria before the latter acquired photosynthetic capabilities (Di Rienzi et al., 2013). Hence FECB32 might be a particularly valuable co-culture to generate new insights into the evolution of and relationship between the phylogenetically closely related *Cyanobacteria* and *Melainabacteria*. In addition, this sample might provide the opportunity to enhance our understanding of the origin of oxygenic photosynthesis and aerobic respiration in *Cyanobacteria*, an area that is currently still poorly understood (Soo et al., 2017).

Interestingly, OTU000109 was also detected in FECB36 and FECB38 (Supplemental Table S2), although at significantly lower abundance (<0.001%). FECB36 and FECB38 were similar to FECB32 in that they were isolated from sites in YNP. Interestingly, FECB32 and FECB38 cluster together (cluster IX) suggesting similar overall microbial community profiles, but separately from FECB36 (Figure 1). The only additional samples that contained OTUs classified as *Melainabacteria*, recruiting >0.1% of the generated reads, were FECB58 and FECB68 with ~0.9% and ~0.2% of their reads to this deeply branched phylum, respectively (Supplemental Table S2). It seems noteworthy that FECB58 and FECB68 were also isolated from hot springs and clustered closely together based on their overall microbiome composition (Clusters V and IV respectively; Figure 1).

### Photosynthetic co-cultures from Hunter’s Hot Spring, Oregon

FECB58 and FECB68 were both isolated from Hunters Hot Spring in Oregon, USA and they shared similar microbial community members. Despite their similar community profile, abundances of the dominant OTUs associated with these two hot spring co-cultures were remarkably different. FECB58 was dominated by 3 OTUs (OTU000014, OTU000024, and OTU000033). OTU000014 was classified as OS-L, an uncultured representative of the phylum *Armatimonadetes*, OTU000024 which was classified as belonging to the *Bacteroidetes* phylum, and OTU000033 which was classified as *Thermosynechococcus*. These OTUs contributed 38%, 29% and 20% of the reads generated from FECB58 respectively. Whereas OTU000014 recruited ~4.9% of all reads generated from FECB68, representing the sixth most abundant OTU in the FECB68 community, OTU000024 and OTU000033 were only present at an abundance <0.0001% in FECB68 (Supplemental Table S2).

FECB68 was dominated by 6 OTUs (i.e. OTU000028, OTU000030, OTU000036, OTU000049, OTU000065, and OTU000014) recruiting ~25.7%, 23.1%, 20.4%, 14.3%, 7.6%, and 4.9% of the reads respectively. OTU000028 was classified as belonging to the genus *Chloroflexus*, whereas OTU000030 and OTU000036 were classified as representative of the genus *Meiothermus* and *Gloeobacter*, respectively. *Chloroflexus* is an anoxygenic phototrophic bacterium that grows at temperatures up to 70°C(Castenholz, 2015) and forms yellow-orange-greenish mats in association with cyanobacteria (Hanada, 2014). Members of the cyanobacterial genus *Gloeobacter* lack thylakoids, and have been proposed to host the earliest ancestors, or a missing link, in the cyanobacteria lineage (Saw et al., 2013). Thus, FECB68 offers a unique opportunity to investigate interspecies interaction between a member of these basal cyanobacteria and the thermophilic phototroph *Chloroflexus*, represented by OTU000028 in this co-culture. As outlined in a recent review by Castenholz (2015), Hunter’s Hot Spring located in Oregon is one of the most studied hot springs in the world and a large repertoire of work has been conducted on this habitat over the last 40 years. However, most of this work was performed prior to the advent of recent molecular and -omics techniques. Hence, sequencing data generated from FECB58 and FECB68 during this study complement previous work and will facilitate new insights into the microbiology of this unique ecosystem.

### Photosynthetic co-cultures from lignocellulosic surfaces with potential to fix nitrogen and degrade aromatic compounds

FECB22 and FECB26 are mesophilic co-cultures collected from similar habitats (i.e. from tree bark and a wooden fence) from two locations (i.e. Hawaii and Bermuda) approximately 9,000 kilometers apart from each other (Supplemental Figure S1 & Table 1). Diversity index calculation placed these two samples in the mid-range of the diversity spectrum of the 26 co-cultures analyzed for this study. The inverse Simpson and Shannon index was calculated at 4.46 and 2.08 for FECB22 and 2.29 and 1.32 for FECB26, respectively (Table 2). Within FECB22, 23 OTUs were identified as individually recruiting more than 0.1% of the generated reads. In contrast, FECB26 contained only 16 OTUs that recruited more than 0.1% of the reads each (Table S2). FECB22, scraped from tree bark in Hawaii, was dominated by 11 OTUs, each recruiting >1% of the reads. The most abundant OTU (OTU000017) was classified as a member of the *Mycoplana*, a genus that contains bacteria capable of aromatic compound degradation (Urakami et al., 1990), and it recruited 40.2% of the reads. OTU000042 (classified as *Rhizobium leguminosarum*), OTU000045 (classified as *Acetobacteraceae*), and OTU000072 (classified as *Cyanobacteria*), were the next most abundant OTUs, recruiting 17.1%, 16.3%, and 5.5% of the reads generated from FECB22 respectively. *Rhizobium leguminosarum* is a well-studied α-proteobacterium capable of N_2_-fixation and “rhizobia” have been suggested repeatedly to facilitate more sustainable agricultural practices through their symbiosis with legumes, reducing the need for nitrogen fertilizer (Marek-Kozaczuk et al., 2017). It remains to be seen if OTU000042 provides N_2_ to the other organisms in this co-culture or if it consumes all of the fixed N_2_ itself. *Acetobacteraceae* are α-proteobacteria often associated with low pH environments and are known for their ability to efficiently synthesize biological cellulose (Rozenberga et al., 2016; Semjonovs et al., 2017). Furthermore, *Acetobacteraceae* have been reported before as some of the dominant players in photosynthetic consortia during soil formation (Mapelli et al., 2011). It would be interesting to explore the agricultural and chemical potential of a minimalistic co-culture composed of the 4 OTUs (i.e. OTU000017, OTU000042, OTU000045 and OTU000072) that dominated FECB22, as they may combine the ability to degrade aromatic compounds and synthesize cellulose while removing nitrogen from the atmosphere. FECB26, on the other hand, was dominated by OTU000010, which recruited 63.2% of the reads generated and it was identified as an unclassified member of the *Nostocales*; a phylogenetic group known for their functional and morphological diversity. Members of the *Sphingomonadaceae* (i.e. OTU000041 and OTU000062), phototropic α-proteobacteria often found in high abundance in environments previously thought to support mostly the growth of cyanobacteria (Tahon and Willems, 2017), contributed to a total of 25.6% of the generated reads. Most interestingly, OTU000017 was also detected within FECB26 recruiting ~1.6% of the reads. It is possible that OTU000017 facilitates a metabolic reaction in which aromatic compounds typically associated with the decomposition of woody material under aerobic conditions are utilized. Further characterization of this organism in co-culture and eventually in axenic culture might provide further clarity if this is the case.

## CONCLUSION

Culture collections can provide easy access to biological samples without the need for extensive resources by the requesting individual, subsequently facilitating new studies and ultimately advancing our understanding of phylogenetic and functional biodiversity. Although some of the diversity of the original microbial community might have been lost due to a cultivation bias, the 16S rRNA based community fingerprints of the 26 photosynthetic co-cultures described here provide a first in-depth glimpse into the taxonomic and functional diversity of communities from extreme environments that were considered for a long time as too harsh to support the growth of complex microbial communities. The extreme conditions that are associated with the habitats from where these co-cultures were collected offer the unique opportunity to study the molecular mechanisms that support the growth of these extremophilic co-cultures and their role in global carbon and nitrogen cycling. Furthermore, an in-depth understanding of these extreme co-cultures holds the potential to discover novel microbial proteins that might render current agricultural, industrial and medical processes more economical and sustainable.

The heterogeneity of the physical parameters reported for the sites where the samples presented in this work were collected, highlights a major challenge (i.e. standardization of protocols) associated with environmental samples and their corresponding metadata (i.e. data describing conditions at each sampling site), specifically when collected during independent sampling efforts. Fortunately, with recent advances in data technologies, the task of data acquisition and dissemination has become less of a challenge. In order to make the best use of these technologies defining a set of minimal information parameters to be recorded during the collection of an environmental sample is of great importance. Similar efforts have been successfully implemented by the Genomic Standards Consortium (GSC) for microbial genomes and metagenomes in the form of the “minimum information about a genome sequence” (MIGS) (Field et al., 2008) and are enforced when describing a novel microbial species (Kampfer et al., 2003).

The identification of **M**inimum **I**nformation about a **C**o-**C**ulture **S**ample (MICCS) would be a significant step in standardizing sample acquisition and maintenance, increasing the value of current and future microbial samples collected from the environment. Developing MICCS and applying them to co-cultures currently available from existing culture depositories is beyond the scope of the work presented here, but we hope that the results presented here will contribute to the initiation of this process and stimulate broad involvement and support from the scientific community and various funding agencies.

## Acknowledgements

This work was funded by the College of Agricultural and Environmental Science and the Microbiology & Biochemistry, Molecular, Cellular, and Developmental Biology Graduate Group at University of California Davis (Davis, CA) and the U.S. Department of Energy (DOE) Joint Genome Institute (JGI) in Walnut Creek, CA. Work conducted by the JGI, a DOE User Facility, is supported by DOE’s Office of Science under Contract No. DE-AC02-05CH11231. We would also thank Drs. Jorge Rodrigues and John Meeks from UC Davis for providing valuable comments and suggestions on how to improve this manuscript.

We would like to dedicate this publication to Professor Dr. Richard Castenholz who passed away during the completion of this work after a long and satisfying journey in the world of Cyanobacteria. He was, and will remain, a great inspiration to many of us.

## Competing Interest Statement

The authors declare no conflicts of interest.

## Author Contributions Statement

Claire Shaw, Charles Brooke, Richard Castenholz, David E. Culley, Matthias Hess, and Susannah G. Tringe wrote the manuscript. Richard Castenholz and Matthias Hess designed the experiment. Erik Hawley and Matthias Hess performed experiment. Michael Barton, David E. Culley, Tijana Glavina del Rio, Miranda Harmon-Smith, Erik Hawley, Matthias Hess, Nicole Shapiro, and Susannah G. Tringe generated the data. Michael Barton, Claire Shaw, Charles Brooke, Morgan P. Connolly, David E. Culley, Javier A. Garcia, Tijana Glavina del Rio, Miranda Harmon-Smith, Erik Hawley, Matthias Hess, and Nicole Shapiro analyzed the data.

## Notes

### Competing Interest Statement

Author Erik Hawley was employed by the company Bayer and Dr, David E. Culley was employed by the company Greenlight Biosciences. The remaining authors declare that the research was conducted in the absence of any commercial or financial relationships that could be construed as a potential conflict of interest.

### Summary of Updates

Title updated

