## Supplementary figures and images for "Phototrophic co-cultures from extreme environments: community structure and potential value for fundamental and applied research"

### Supplemental Figure 1

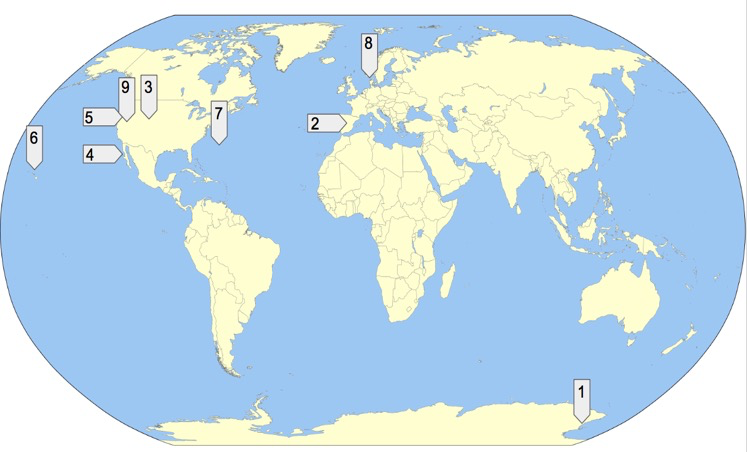

### Supplemental Figure 2

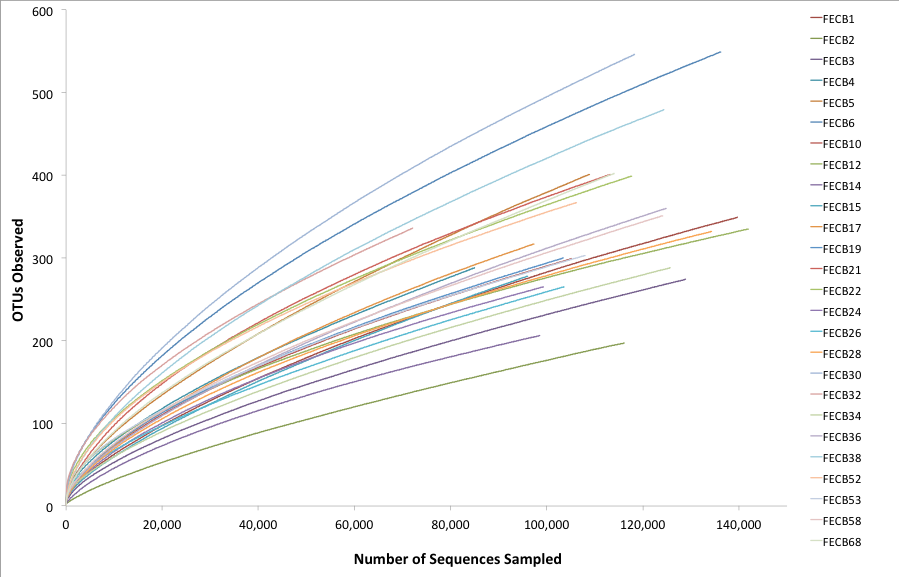
